# Short ‘1.2× genome’ infectious clone initiates deltavirus replication in *Boa constrictor* cells

**DOI:** 10.1101/2021.10.01.462842

**Authors:** Leonora Szirovicza, Udo Hetzel, Anja Kipar, Jussi Hepojoki

**Affiliations:** University of Helsinki, Medicum, Department of Virology, Helsinki, Finland; Institute of Veterinary Pathology, Vetsuisse Faculty, University of Zürich, Zürich, Switzerland; Department of Veterinary Biosciences, Faculty of Veterinary Medicine, University of Helsinki, Helsinki, Finland

**Author notes:** Corresponding author* Correspondence should be addressed to Leonora Szirovicza, *Mailing address:* University of Helsinki, Department of Virology, P.O. Box 21, Haartmaninkatu 3, FI-00014 University of Helsinki, Finland.

## Abstract

Human hepatitis D virus (HDV), discovered in 1977, represented the sole known deltavirus for decades. The dependence on hepatitis B virus (HBV) co-infection and its glycoproteins for infectious particle formation led to the assumption that deltaviruses are human-only pathogens. However, since 2018, several reports have described identification of HDV-like agents from various hosts but without co-infecting hepadnaviruses. Indeed, we demonstrated that Swiss snake colony virus 1 (SwSCV-1) uses arenaviruses as the helper for infectious particle formation, thus shaking the dogmatic alliance with hepadnaviruses for completing deltavirus life cycle. *In vitro* systems enabling helper virus-independent replication are key for studying the newly discovered deltaviruses. Others and we have successfully used constructs containing multimers of the deltavirus genome for the replication of various deltaviruses via transfection in cell culture. Here, we report the establishment of deltavirus infectious clones with 1.2× genome inserts bearing two copies of the genomic and antigenomic ribozymes. We used SwSCV-1 as the model to compare the ability of the previously reported “2× genome” and the “1.2× genome” plasmid constructs/infectious clones to initiate replication in cell culture. Using immunofluorescence, qRT-PCR, immuno- and northern blotting, we found the 2× and 1.2× genome clones to similarly initiate deltavirus replication *in vitro* and both induced a persistent infection of snake cells. We hypothesize that duplicating the ribozymes facilitates the cleavage of genome multimers into unit-length pieces during the initial round of replication. The 1.2× genome constructs enable easier introduction of modifications required for studying deltavirus replication and cellular interactions.

**IMPORTANCE:** Hepatitis D virus (HDV) is a satellite virus infecting humans with strict association to hepatitis B virus (HBV) co-infection because HBV glycoproteins can mediate infectious HDV particle formation. For decades, HDV was the sole representative of deltaviruses, which had led to hypotheses suggesting that it evolved in humans, the only known natural host. Recent sequencing studies have led to the discovery of HDV-like sequences across a wide range of species, representing a paradigm shift in deltavirus evolution. Molecular biology tools such as infectious clones, which enable initiation of deltavirus infection without helper virus, are key to demonstrate that the recently found deltaviruses are capable of independent replication. Such tools will enable identification of the potential helper viruses. Here, we report a 1.2× genome copy strategy for designing plasmid-based infectious clones to study deltaviruses and to demonstrate that plasmid delivery into cultured snake cells sufficiently initiates replication of different deltaviruses.

## INTRODUCTION

Hepatitis D virus (HDV) is a unique human pathogen. Three years after its discovery in 1977 in liver specimens of chronically hepatitis B (HBV) infected patients (1), Rizzetto and colleagues identified it as a satellite virus of HBV (2). One can contract HDV in two different ways, either through acute co-infection with HBV or through superinfection as a chronic HBV carrier. HBV and HDV co-infection is clinically more severe than HBV mono-infection; however, the infection usually resolves resulting in the clearance of both viruses. Superinfection of a chronic HBV carrier by HDV results in the most severe form of viral hepatitis; these patients often face hepatic cirrhosis and development of hepatocellular carcinoma (3). HDV is a satellite virus that utilizes the envelope proteins of HBV to assemble infectious viral particles; however, the replication of HDV within the host cell proceeds independently of HBV (4). The single-stranded RNA genome of HDV is around 1.7 kilonucleotides long although because of high self-complementary, it forms a double-stranded rod-like structure (5, 6). Within the cell, HDV gives rise to three different RNA species: the genome, the antigenome, and the mRNA. The antigenome is the exact complement of the genome, while the mRNA mediates the expression of the delta antigen (DAg), the sole protein encoded by the HDV genome (5, 7). During the viral life cycle, the DAg is present in two different forms, small (SDAg) and large DAg (LDAg) (8). Cellular editing mediated by adenine deaminase 1 (ADAR1) converts the amber stop codon of the SDAg on the antigenomic strand to a tryptophan codon, thus allowing the extension of the protein by 19 additional amino acids (9, 10). The two forms of the protein not only differ in their length but they also have vastly different roles. The SDAg promotes viral replication, while the LDAg inhibits it and shifts the viral life cycle towards packaging (8). The host cell’s RNA polymerases mediate the replication of the HDV genome, which occurs via a double rolling circle mechanism (11). A curious feature of HDV is the presence of ribozyme sequences in both its genome and antigenome (12). The ribozymes cut the multimeric HDV RNA species produced during the rolling circle replication into unit-length pieces (11).

HDV was the sole representative of the unassigned genus *Deltavirus* until 2018 (13). The discovery of HDV-like sequences in birds and snakes in 2018 marked the beginning of a new chapter in deltavirus research by broadening the potential host spectrum (14, 15). First, an HDV-like sequence was discovered in waterfowl during a meta-transcriptomic study without traces of HBV or hepadnaviral reads but influenza A virus reads instead (14). Co-incidentally, we reported the identification of a deltavirus in *Boa constrictors* (Swiss snake colony virus 1, SwSCV-1, initially known as snake deltavirus, SDeV) in co-infection with reptarena- and hartmaniviruses but in the absence of hepadnaviral reads (15). The next paradigm shift of deltavirus research was the broadening of the scope of putative HDV helper viruses to include vesiculo-, flavi- and hepaciviruses (16). The researchers further showed that HDV forms infectious particles using the glycoproteins (GPs) of the aforementioned viruses, not only in liver, but also in kidney cells. Soon after, we managed to isolate SwSCV-1 along with co-infecting reptarena- and hartmaniviruses from brain homogenates of an infected snake (17). We utilized persistently SwSCV-1 infected cell cultures to demonstrate that reptarena- or hartmanivirus superinfection results in the egress of infectious SwSCV-1 particles, adding to the evidence that reptarena- and hartmaniviruses are the likely helpers of the virus (17). We further demonstrated that expression of arena- and orthohantavirus GPs in the persistently infected cultures also induces infectious particle formation (17). The aforementioned reports provoked several meta-transcriptomic studies, resulting initially in the identification of deltaviruses in subterranean termites, fish, Asiatic toad, and Chinese fire belly newt in 2019 (18). In 2020, Paraskevopoulou and colleagues found a deltavirus in Tome’s spiny rats without sequences for a helper virus, but including evidence for autonomous replication in cell culture and the host (19). Metatranscriptomic studies revealed more deltaviruses in common vampire bats, a lesser dog-like bat, white-tailed deer, eastern woodchuck, passerine birds, zebra finch, lantern fish, and amboli leaping frogs (20-22). The recent findings have sparked deltavirus research and raised questions about the evolutionary origin of deltaviruses (23-25). The identification of deltaviruses across several taxa led to the establishment of a new realm, *Ribozyviria*, with family *Kolmioviridae* including eight genera (26, 27).

A number of tools to initiate HDV replication *in vitro* and to mimic infection have been developed over the years. The first and probably most widely used method is the transfection of cells with a plasmid containing a trimer of the entire HDV genome under the control of the simian virus 40 promoter (4). In theory, the transfection results in multimeric RNA transcripts of the HDV genome, which resembles the rolling circle replication of the virus during actual infection. It is also possible to initiate the replication by transfecting a vector carrying a monomer of the HDV genome, but in this case, an HDAg expressing plasmid needs to be provided *in trans* (28). Macnaughton and Lai showed that direct RNA transfection with HDAg provided *in trans* can overcome the use of artificial DNA intermediates to initiate replication. Interestingly, they found that transfection with 1.2× genome-length RNA resulted in the most efficient replication, perhaps because the ribozymes at both ends of the genome were in duplicate (29). In our previous study, we constructed a plasmid containing the entire SwSCV-1 genome in duplicate in head-to-tail fashion (17), following the idea of the trimeric HDV constructs (4). The transfection of the “2×” construct initiates efficient SwSCV-1 replication in snake cells, eventually leading to persistent infection (17). The ‘genome-dimer or 2×’ plasmid principle was also successfully adapted for Tome’s spiny rat virus 1 (TSRV-1), *Taeniopygia guttata* deltavirus, and *Marmota monax* deltavirus (19, 21). Here, we describe a novel construct containing 1.2× SwSCV-1 genome, which in a manner similar to the previously described 2× SwSCV-1 infectious clone (17), initiates virus replication, produces infectious particles upon superinfection with Haartman institute snake virus 1 (HISV-1) and results in persistent infection of the cells. We based our 1.2× infectious clone on the RNA transfection studies of Macnaughton and Lai (29) with the aim to generate shorter DNA constructs to facilitate introduction of mutations or modifications to the virus genome. Additionally, we wanted to eliminate the T7 promoter we used in the 2× SwSCV-1 infectious clone to avoid the risk of DAg expression via this promoter due to cellular polymerases, which is reported to occur in mammalian cells (30, 31).

## MATERIALS AND METHODS

### Cell culture and superinfection

The study made use of previously described cultured *Boa constrictor* kidney cells, I/1Ki, (32) and persistently SwSCV-1 infected I/1Ki cells, I/1Ki-Δ (17). The cells were maintained in Minimal Essential Medium Eagle (Sigma Aldrich) supplemented with 10% fetal bovine serum (Gibco), 200 mM L-glutamine (Sigma Aldrich), 100 μg/ml of streptomycin (Sigma Aldrich), and 100 U/ml of penicillin (Sigma Aldrich) in an incubator at 30°C with 5% CO_2_.

To establish another persistently SwSCV-1 infected I/1Ki cell line, designated I/1Ki-1.2×Δ, we transfected I/1Ki cells with a plasmid containing 1.2 copies of the SwSCV-1 genome (described below) and maintained the cells as described above and earlier (17).

To study infectious particle formation of the SwSCV-1 infected cell lines, we conducted superinfection studies with HISV-1 (33), earlier demonstrated to efficiently serve as the helper for SwSCV-1 (17). The superinfection studies and detection followed the protocol described (17).

### Plasmids and cloning

We ordered synthetic genes from Gene Universal for the 1.2× copies of the following deltaviruses: SwSCV-1 (initially known as snake deltavirus, GenBank accession number: NC_040729.1)(17), Tome’s spiny rat virus 1 (TSRV-1, initially known as rodent deltavirus, MK598005.2)(19), dabbling duck virus 1 (DabDV-1, initially known as avian HDV-like agent, NC_040845.1)(14), Chusan Island toad virus 1 (CITV-1, initially known as toad HDV-like agent, MK962760.1)(18), and HDV-1 (M21012.1). Each synthetic gene, flanked by EcoRV restriction sites, contained the full genome with additional nucleotides to include both the genomic and antigenomic ribozymes twice (Figure 1). Subcloning of the constructs into pCAGGS followed the procedures described in Szirovicza et al, 2020 (17). Briefly, FastDigest EcoRV (ThermoFisher Scientific) served to restriction digest the inserts, followed by purification after agarose gel electrophoresis using the GeneJET Gel extraction kit (ThermoFisher Scientific). T4 DNA ligase (ThermoFisher Scientific) served to ligate the purified inserts into pCAGGS/MCS plasmid (34) purified from agarose gel by the GeneJET Gel extraction kit (ThermoFisher Scientific) after linearization with Fast Digest EcoRI and XhoI (ThermoFisher Scientific) restriction enzymes and T4 DNA polymerase (ThermoFisher Scientific) blunting. We plated chemically competent *E. coli* (DH5α strain) transformed with the ligation products on Luria-Broth (LB) agar plates with 100 μg/ml of ampicillin, and incubated overnight (O/N) at 37°C. We picked single colonies and transferred them into 5 ml of LB medium (10 g/l tryptone, 10 g/l NaCl, 5 g/l yeast extract), followed by O/N incubation at 37°C (220 rpm), after which the GeneJET Plasmid Miniprep Kit (ThermoFisher Scientific) served for plasmid isolation from 2 ml of the O/N culture. The DNA Sequencing and Genomic Laboratory, Institute of Biotechnology, University of Helsinki performed Sanger sequencing of the preparations and confirmed these to contain an insert of the correct size. For each virus, we selected two clones, one each with the insert in genomic and in antigenomic orientation for plasmid stock preparation using ZymoPURE II Plasmid Maxiprep Kit (Zymo Research).

**Figure 1.**
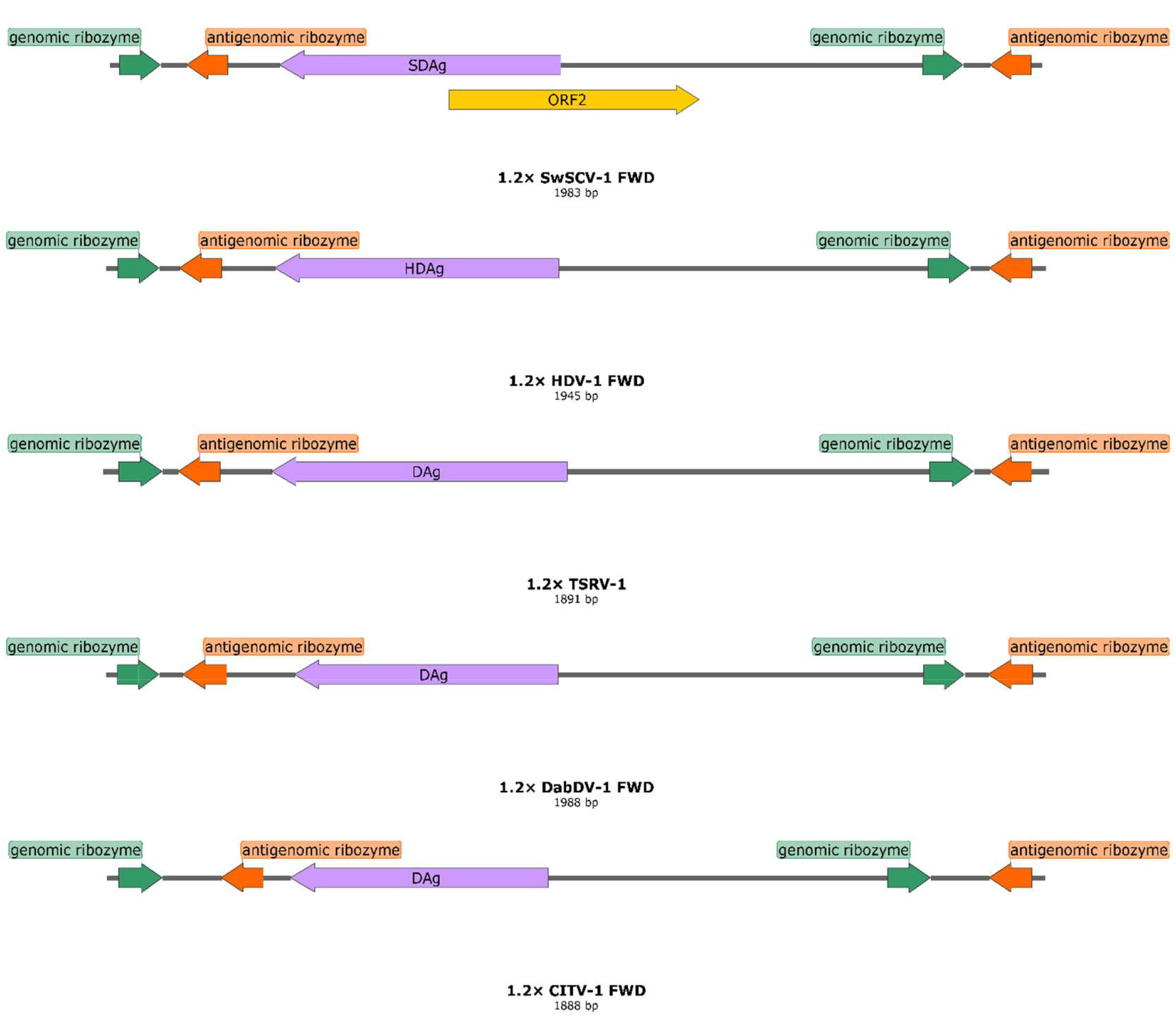
Schematic representation of the “1.2× genome” deltavirus inserts in forward (FWD)/genomic orientation. We cloned the 1.2× genome of the displayed deltaviruses: swiss snake colony virus 1 (SwSCV-1, GenBank accession: NC_040729.1, 1.15× genome), human hepatitis D virus genotype 1 (HDV-1, M21012.1, 1.16× genome), Tome’s spiny rat virus 1 (TSRV-1, MK598005.2, 1.13× genome), Dabbling duck virus 1 (DabDV-1, NC_040845.1, 1.17× genome) and Chusan Island toad virus 1 (CITV-1, MK962760.1, 1.22× genome) into pCAGGS/MCS plasmid, both in genomic (FWD – shown in this figure) and antigenomic (REV) orientation. Each of the inserts, approximately 1.2× of the genome size, contain a single copy of the genomes flanked from each end by both the genomic and antigenomic ribozymes. The images were created using SnapGene Viewer.

### Transfection

Lipofectamine 2000 (ThermoFisher Scientific) reagent served for transfection of I/1Ki cells as described (17, 35). Briefly, we mixed 500 ng of plasmid DNA in 50 μl of OptiMEM (ThermoFisher Scientific) and 3 μl of Lipofectamine 2000 in 47 μl of OptiMEM (ThermoFisher Scientific) by pipetting up and down, and allowed the complexes to form for 15-30 minutes at room temperature (RT). We added 1 ml of trypsinized cells (suspension containing approximately 1.8 cm^2^ of cells per ml) to the mixture and allowed the suspension stand at RT for 15-30 min before plating. At 5-6 h post plating, we replaced the transfection mixture by fully supplemented medium and incubated the cells as described above. We scaled up the above reaction volumes depending on the amount of cells needed for each experiment.

### Western blot (WB)

For WB, we washed the cells grown on plates or flasks twice with PBS, scraped them into PBS, pelleted by centrifugation (500 × g, 3-5 min), lysed the cell pellets by RIPA buffer (50 mM Tris, 150 mM NaCl, 1% Tx-100, 0.1% SDS, 0.5% sodium deoxycholate, protease inhibitor cocktail), and measured the protein concentration using the Pierce^™^ BCA Protein Assay Kit (ThermoFisher Scientific). For comparing DAgs of different deltaviruses, we collected from a 12-well plate non-transfected I/1Ki cells and those transfected with FWD and REV constructs at 4 days post transfection in 100 μl of Laemmli sample buffer after two washes with PBS. We separated an equal amount of protein (or volume, 30 μl/lane, for the experiments done on 12-well plate) for each sample on SDS-PAGE using 4–20% Mini-PROTEAN® TGX gels (Bio-Rad), and immunoblotted as described (36). We used 1:4000 dilution for the rabbit anti-SDAg primary antibody (15) or 1 μg/ml of IgG affinity purified from anti-SwSCV-1 rabbit antiserum (17) using recombinant human DAg as described (17), 1:10,000 for AlexaFluor 680 donkey anti-mouse (IgG) (ThermoFisher Scientific), and 1:10,000 for IRDye 800CW donkey anti-rabbit (IgG) (LI-COR Biosciences). The Lab Vision^™^ pan-actin mouse monoclonal antibody (ThermoFisher Scientific) used at a 1:200 dilution served for detection of β-actin. The Odyssey Infrared Imaging System (LI-COR Biosciences) was employed to record the results.

### Immunofluorescence staining

For immunofluorescence (IF) staining, we plated the cells on collagen coated (10 μg/cm^2^ type I rat tail collagen [BD Biosciences] in 25 mM acetic acid, O/N at 4°C) on CellCarrier-96 Ultra plates (PerkinElmer). After the removal of culture media, cells were fixed by incubation in 4% paraformaldehyde in PBS for ∼15 min at RT. The IF staining followed the protocol described (17). We used directly labelled anti-SDAg-AF488 (17) at 1:500 dilution, rabbit anti-SDAg antiserum at 1:4,000 dilution, and 1:1,000 of either Alexa Fluor 488- or 594-labeled donkey anti-rabbit immunoglobulin (ThermoFisher Scientific) as the secondary antibody. The Opera Phenix High Content Screening System (PerkinElmer), provided by FIMM (Institute for Molecular Medicine Finland) High Content Imaging and Analysis (FIMM-HCA), served for imaging of the plates stored in the dark at 4°C.

### Quantitative reverse transcription PCR (qRT-PCR)

qRT-PCR served for quantification of viral RNA in the cells. The primers and probe were the following: forward primer 5’-GAAAGACGCGACAACTGTGAGTC-3’, reverse primer 5’-GTCTAGTCCCGTTCCGGTTCTATG-3’, and probe 5’ 6-Fam (carboxyfluorescein)-GGAGATCCGAGAGGGGAGAAGAGGAGAGGTC-BHQ (black hole quencher)-1 3’, which target SwSCV-1 RNA in genomic orientation. We isolated RNA for qRT-PCR using the GeneJET RNA Purification Kit (ThermoFisher Scientific) with the addition of carrier RNA when purifying RNA from cell culture supernatants. We used TaqMan® Fast Virus 1-Step Master Mix (ThermoFisher Scientific) to set up 10 μl (half volume) reactions according to the manufacturer’s recommendations with the addition of 8% DMSO to prevent secondary structure formation. The AriaMX real-time PCR system (Agilent) served for thermal cycling of the duplicate samples with the recommended conditions: reverse transcription for 5 min at 50°C; initial denaturation for 20 s at 95°C; and two amplification steps at 95°C for 3 s and 60°C for 30 s repeated for 40 cycles.

To generate a control RNA for copy-level quantification, we used the SwSCV-1-FWD plasmid described in Szirovicza et al., 2020 (17). Briefly, FastDigest SmaI (ThermoFisher Scientific) following manufacturer’s protocol served for linearization of the plasmid. The GeneJet Gel Purification Kit (ThermoFisher Scientific) served for purification of the linearized plasmid after agarose gel separation. We used the TranscriptAid T7 High Yield Transcription Kit (ThermoFisher Scientific) according to the manufacturer’s protocol to *in vitro* transcribe the target RNA. Subsequently, we purified the RNA by the GeneJET RNA Purification Kit (ThermoFisher Scientific), diluted it into diethyl pyrocarbonate (DEPC) treated water, and stored the RNA in aliquots at -80°C until use. The NanoDrop 2000 spectrophotometer (ThermoFisher Scientific) served for quantification of the control RNA and an online copy number calculator (http://endmemo.com/bio/dnacopynum.php) for converting the concentration to RNA copies per microliter. We ran the control RNA as 10-fold dilution series in duplicates for each run to generate a standard curve for estimation of RNA copy numbers in cell and cell culture supernatant samples.

To normalize the SwSCV-1 RNA levels against a house-keeping gene, we ordered the following primers and probe for the detection of *Boa constrictor* glyceraldehyde-3-phosphate dehydrogenase (GAPDH): forward primer: 5’ CTGGTATGACAACGAATA 3’, reverse primer: 5’ CAGTCTTTACTCCTTAGATG 3’, and probe 5’ 6-Fam (carboxyfluorescein)-TGAACCAACAAGTCTACCACACG -BHQ-1 3’. Reference assembly using *Python bivitattus* GAPDH (GenBank accession: XM_007429612.3) as the template in Unipro UGENE (37) served to obtain the mRNA for *B. constrictor* GAPDH from the reads of our earlier metatranscriptomic studies (33, 38-40).

### Near-infrared fluorescent northern blot

For northern blot analysis, TRIzol^™^ reagent (ThermoFisher Scientific) used according to the manufacturer’s recommendations served for isolating RNA from either T75 or T175 flasks of I/1Ki, I/1Ki-2×Δ and I/1Ki-1.2×Δ. The NanoDrop 2000 spectrophotometer (ThermoFisher Scientific) was employed for quantification of the RNA solubilized into formamide after 1:100 dilution in DEPC-treated water. Subsequently, we ran 3-5 μg of RNA to detect genomic SwSCV-1 RNA, 20-25 μg to detect antigenomic RNA, and an ssRNA ladder (New England Biolabs) on agarose-formaldehyde gels using the tricine/triethanolamine buffer system and method described by Mansour and Pestov (41). Then, we transferred the RNAs from the gel to HybondTM-N+ nylon membrane (GE Healthcare) by capillary transfer O/N and cross-linked the RNA to the membrane by an ultraviolet cross-linker (120 mJ/cm^2^ at 254 nm) (Analytik Jena). Incubation in prehybridization buffer (5× sodium saline citrate buffer [SSC], 5× Denhardt’s solution [ThermoFisher Scientific], 1% SDS, 50% formamide and 100 μg ultrapure herring sperm DNA [ThermoFisher Scientific]) at 68°C or 48°C (depending on the probe) for 2-4 h served for blocking the non-specific binding sites on the membrane. The following probes (all from Metabion International Ag): ladder-targeting (5’-IR800-AAGCAGGTCCTCGTCGCCGTACACCTCATCATACA-3’), SwSCV-1 genome-targeting (5’-IR800-GCTCTCCCCGGCAAGTCTTCTATTTCTGTCCTTCC-3’), SwSCV-1 antigenome-targeting (5’-IR800-GGAAGGACAGAAATAGAAGACTTGCCGGGGAGAGC-3’), or SDAg mRNA-targeting (5’-IR800-TAATCTCTTTCGGTGGTAGCCTCGAGCCGCCATCC-3’) diluted in prehybridization buffer served for detection. We performed the hybridization O/N at 68°C (using the genome targeting probe) or 48°C (using the antigenome and mRNA targeting probes), and washed the membrane once with 2× SSC and 0.1% SDS for 10 min at 68°C/48°C, followed by two 10 min washes with 1× SSC and 0.1% SDS at 68°C/48°C. After the washes, the Odyssey Infrared Imaging System (LI-COR Biosciences) served to record the results. The protocol was adapted from Miller et al., 2018 (42).

## RESULTS

### Transfection with 1.2× genome constructs initiates deltavirus replication

In our previous study, we showed in a transfection-based assay that a plasmid bearing the SwSCV-1 genome in duplicate could initiate SwSCV-1 replication in cell culture with highest efficacy observed in boid kidney cells (17). We hypothesized, based on RNA transfection studies with HDV (11), that a plasmid bearing a shorter 1.2× genome-length insert would suffice to initiate translation. With the idea that duplicating the antigenomic and genomic ribozymes would better facilitate replication, we ordered synthetic genes representing the 1.2× genome of SwSCV-1, TSRV-1, DabDV-1, CITV-1 and HDV-1. Figure 1 shows the organization of the synthetic blocks that we subsequently cloned in both reverse and forward orientation into a pCAGGS/MCS expression vector under the CAG promoter. Unlike in our earlier study, in which we included an additional T7 promoter in the antigenomic orientation, upstream of the DAg to the synthetic construct (17), we did not include additional promoters that could unintentionally facilitate DAg translation. The resulting constructs we named according to the 1.2× “deltavirus” FWD and 1.2× “deltavirus” REV scheme. The FWD constructs drive transcription of the respective deltavirus antigenome, due to which the expression or translation of SDAg should only occur following virus replication. On the other hand, the REV constructs would generate antigenomic transcripts that would likely also mediate the SDAg translation. To test if the shorter constructs will facilitate replication, and to estimate the cross-reactivity of our rabbit anti-SwSCV-1 DAg antiserum, we transfected I/1Ki cells with each of the constructs. IF staining of the cells transfected with REV constructs for DAg at 4 days post transfection (dpt) showed that the anti-SwSCV-1 DAg antiserum clearly cross-reacted with TSRV-1 and HDV-1, but also to some extent with DabDV-1 and CITV-1 (Figure 2A). We detected DAg in cells transfected with the FWD constructs of HDV-1, TSRV-1, and SwSCV-1, suggesting that transfection resulted in replication initiation (Figure 2B). The staining for DAg of DabDV and CITV-1 was much less prominent on the cells transfected with FWD construct; however, the results suggest that replication occurs also with these viruses. We also performed WB on the transfected cells and we were able to detect DAgs of SwSCV-1, HDV-1, TSRV-1 and DabDV-1 4 days after transfection with the REV constructs. However, after transfection with the FWD constructs, we were only able to detect bands for DAgs of SwSCV-1, HDV-1 and TSRV-1 (Figure 2C). The observed differences between the results of IF staining and WB could be due to the ability of the antibody to bind conformational epitopes in case of IF staining, however, in WB this is not possible.

**Figure 2.**
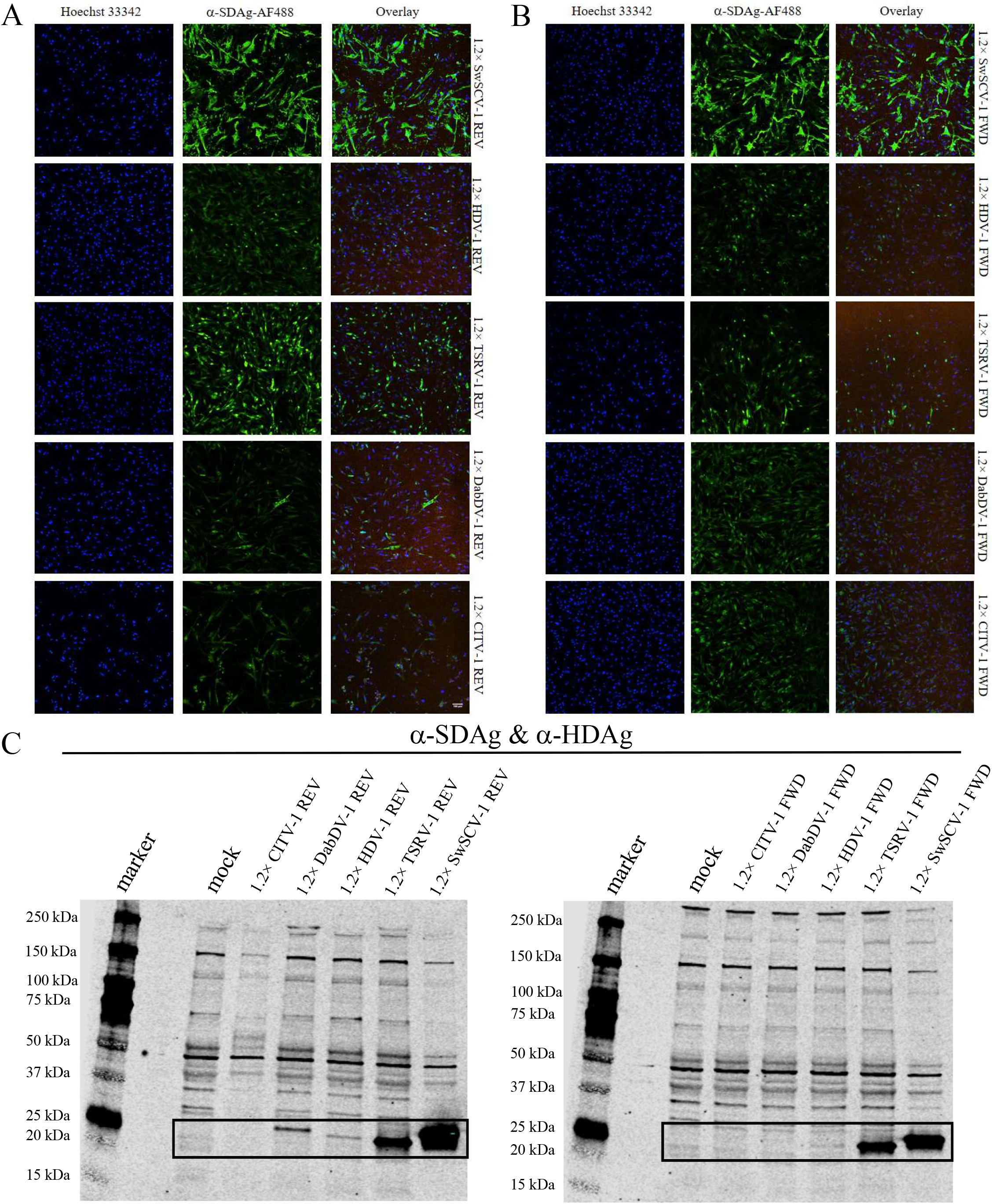
SwSCV-1 DAg antiserum cross-reactivity with the DAg of different deltaviruses. **A)** I/1Ki cells transfected with 1.2× SwSCV-1, HDV-1, TSRV-1, DabDV-1 and CITV-1 REV constructs were stained for the DAg at 4 days post transfection using rabbit α-SwSCV-1 DAg antiserum. **B)** I/1Ki cells transfected with 1.2× SwSCV-1, HDV-1, TSRV-1, DabDV-1 and CITV-1 FWD constructs and stained for the DAg 4 days post transfection using rabbit α-SwSCV-1 DAg antiserum. Hoechst 33342 served for detection of the nuclei (left panels) and AlexaFluor 488 labelled donkey anti-rabbit IgG as the secondary antibody for DAg detection (middle panels). The right panels show overlay of the nuclear and DAg staining. The images were captured using Opera Phenix High Content Screening System (PerkinElmer) with 20× objective. **C)** I/1Ki cells transfected with 1.2× SwSCV-1, HDV-1, TSRV-1, DabDV-1 and CITV-1 REV constructs (left panel) and FWD constructs (right panel) were submitted for western blot at 4 days post transfection. The samples were separated on 4– 20% Mini-PROTEAN TGX gels (Bio-Rad), transferred onto nitrocellulose, and the membranes were probed with rabbit α-SwSCV-1 DAg antiserum and affinity purified α-HDAg antibody. We loaded 1/3 volume of the 1.2× SwSCV-1 REV and FWD samples. The bands corresponding to the different DAgs are marked with the black rectangle. The results were recorded using Odyssey Infrared Imaging System (LI-COR Biosciences).

### The 1.2× and 2× SwSCV-1 genome infectious clones induce similar infection as judged by antigen expression and replication

To compare the ability of the shorter 1.2× genome length constructs to initiate replication, we transfected I/1Ki cells with 1.2× SwSCV-1 FWD and 1.2× SwSCV-1 REV and compared the antigen expression and SwSCV-1 RNA production between 1 and 5 dpt. IF staining of the transfected cells at 1-4 dpt demonstrates that both 1.2× and 2× genome constructs efficiently drive the expression of DAg (Figure 3). To better estimate the amount of DAg produced by the different constructs, we performed western blotting on samples collected between 1 and 4 dpt. The results showed that similarly to the 2× genome constructs, the transfection of 1.2× constructs led to an increasing amount of SDAg expression during the interval studied (Figure 4A). As the amount of DAg appeared to increase until 4 dpt and because the results of the IF staining suggested minute differences in the amount of DAg expression through the different construct, we performed WB analysis of samples collected at 5 dpt. The results indicate that the SDAg expression level in the cells is similar at 5 dpt, regardless of the construct used for transfection (Figure 4B). DAg expression in REV constructs could be driven by the CAG promoter of the plasmid, but since levels were similar at 5 dpt we interpret the result to suggest that DAg expression is due to replication. To add another dimension to the determination of the replication efficiency, we analyzed I/1Ki cells transfected with the 4 different constructs for SwSCV-1 RNA levels at 3 and 6 dpt by qRT-PCR. The 2× SwSCV-1 genome construct appeared to initiate replication more slowly as demonstrated by the lower amount of SwSCV-1 RNA at 3 dpt (Figure 4C). However, at 6 dpt the cells transfected with each of the constructs showed similar SwSCV-1 RNA levels when normalized against GAPDH mRNA (Figure 4C), supporting the WB-based interpretation that all of the studied constructs can initiate replication.

**Figure 3.**
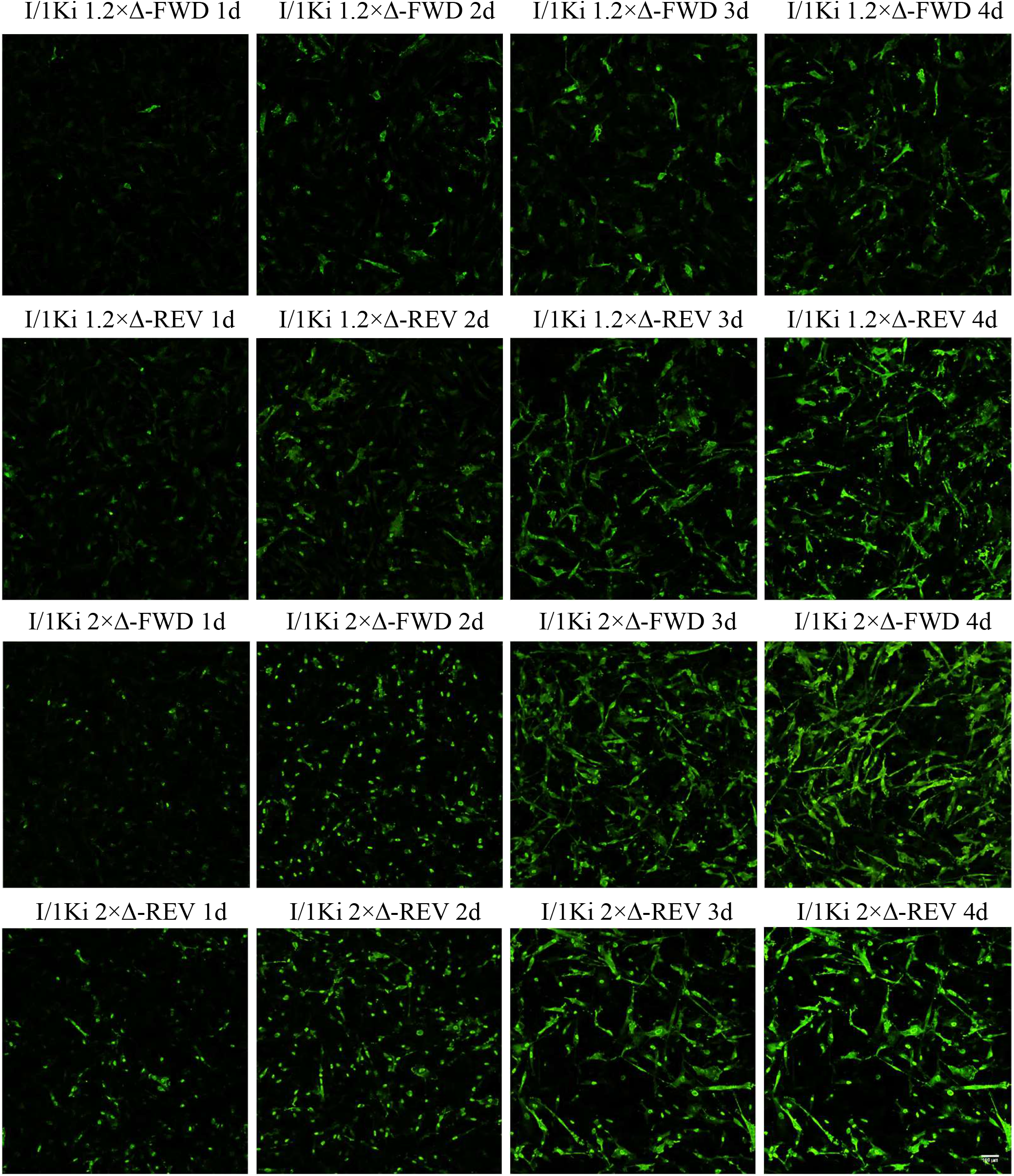
Expression of DAg in I/1Ki cells following transfection with 2× and 1.2× genome SwSCV-1 FWD and REV plasmids. I/1Ki cells transfected with 2× and 1.2× SwSCV-1 FWD and REV plasmids were fixed and stained for the DAg using rabbit α-SwSCV-1 DAg antiserum at 1-4 days post transfection. AlexaFluor 488 labelled donkey anti-rabbit IgG served as the secondary antibody for DAg detection. The images were captured using Opera Phenix High Content Screening System (PerkinElmer) with 20× objective.

**Figure 4.**
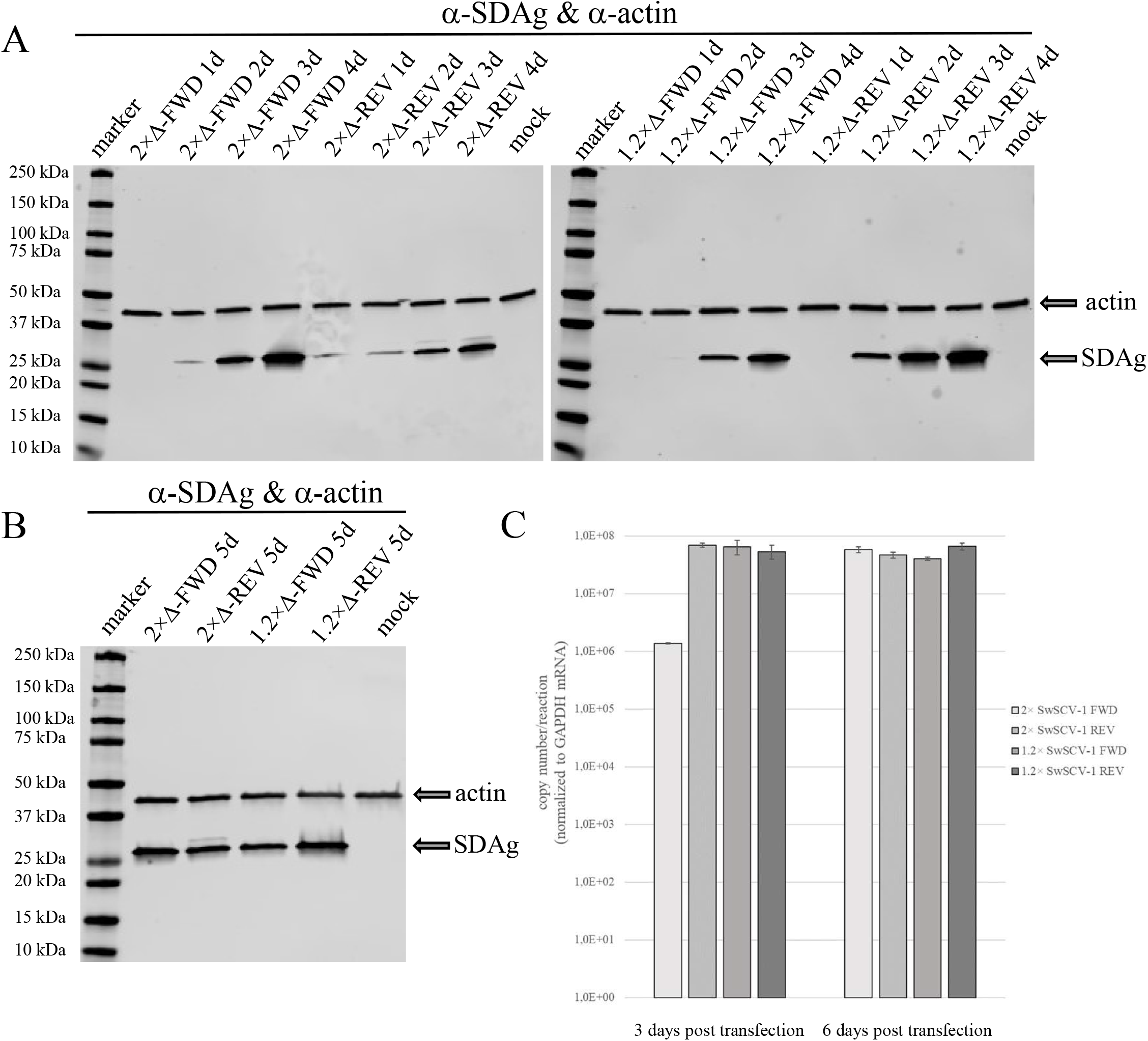
Western blot of I/1Ki cells after transfection with 2× and 1.2× SwSCV-1 (2×Δ and 1.2×Δ respectively) FWD and REV constructs. **A)** Samples of I/1Ki cells transfected with 2×Δ-FWD, 2×Δ-REV, 1.2×Δ-FWD, and 1.2×Δ-REV constructs collected at 1-4 days post transfection were separated on 4–20% Mini-PROTEAN TGX gels (Bio-Rad), transferred onto nitrocellulose, and the membranes were probed with rabbit α-SwSCV-1 DAg antiserum and mouse monoclonal anti-pan actin antibody. The left panel shows 2× and the right panel 1.2× genome constructs. The results were recorded using Odyssey Infrared Imaging System (LI-COR Biosciences). **B)** Samples of I/1Ki cells transfected with 2×Δ-FWD, 2×Δ-REV, 1.2×Δ-FWD, and 1.2×Δ-REV constructs collected at 5 days, analyzed as described in **A. C)** RNA isolated from I/1Ki cells transfected with 2×Δ-FWD, 2×Δ-REV, 1.2×Δ-FWD, and 1.2×Δ-REV constructs at 3 and 6 days post transfection were analyzed by qRT-PCR targeting genomic SwSCV-1 RNA. *In vitro* transcribed RNA target served for obtaining a standard curve to convert cycle threshold values into copy numbers. qRT-PCR targeting GAPDH mRNA served for normalizing the results between samples. The y-axis shows copy numbers/reaction. The error bars represent standard deviation.

### Superinfection of cells transfected with 1.2× SwSCV-1 FWD construct induces infectious particle formation

To show that the 1.2× SwSCV-1 construct not only initiates virus replication in cell culture, but also induces infectious particle formation in the presence of a suitable helper virus, we superinfected 1.2× and 2× SwSCV-1 FWD transfected cells with HISV-1, a hartmanivirus demonstrated to act as a helper for SwSCV-1 (17). We titrated the supernatants collected at 3, 6 and 9 dpi with HISV-1 on clean I/1Ki cells and used supernatants collected from non-superinfected cells as the control. IF staining of cells inoculated with the supernatants at 4 dpi for SDAg served for detecting the infected cells (Figure 5A). We determined the number of infectious units by counting the fluorescent foci at each time point and the results showed I/1Ki cells transfected with 1.2× or 2× SwSCV-1 FWD constructs to be equally effective in producing infectious particles following superinfection (Figure 5B). As observed for 2× SwSCV-1 FWD in our earlier study (17), the non-superinfected 1.2× SwSCV-1 FWD cells were not able to produce infectious SwSCV-1 particles.

**Figure 5.**
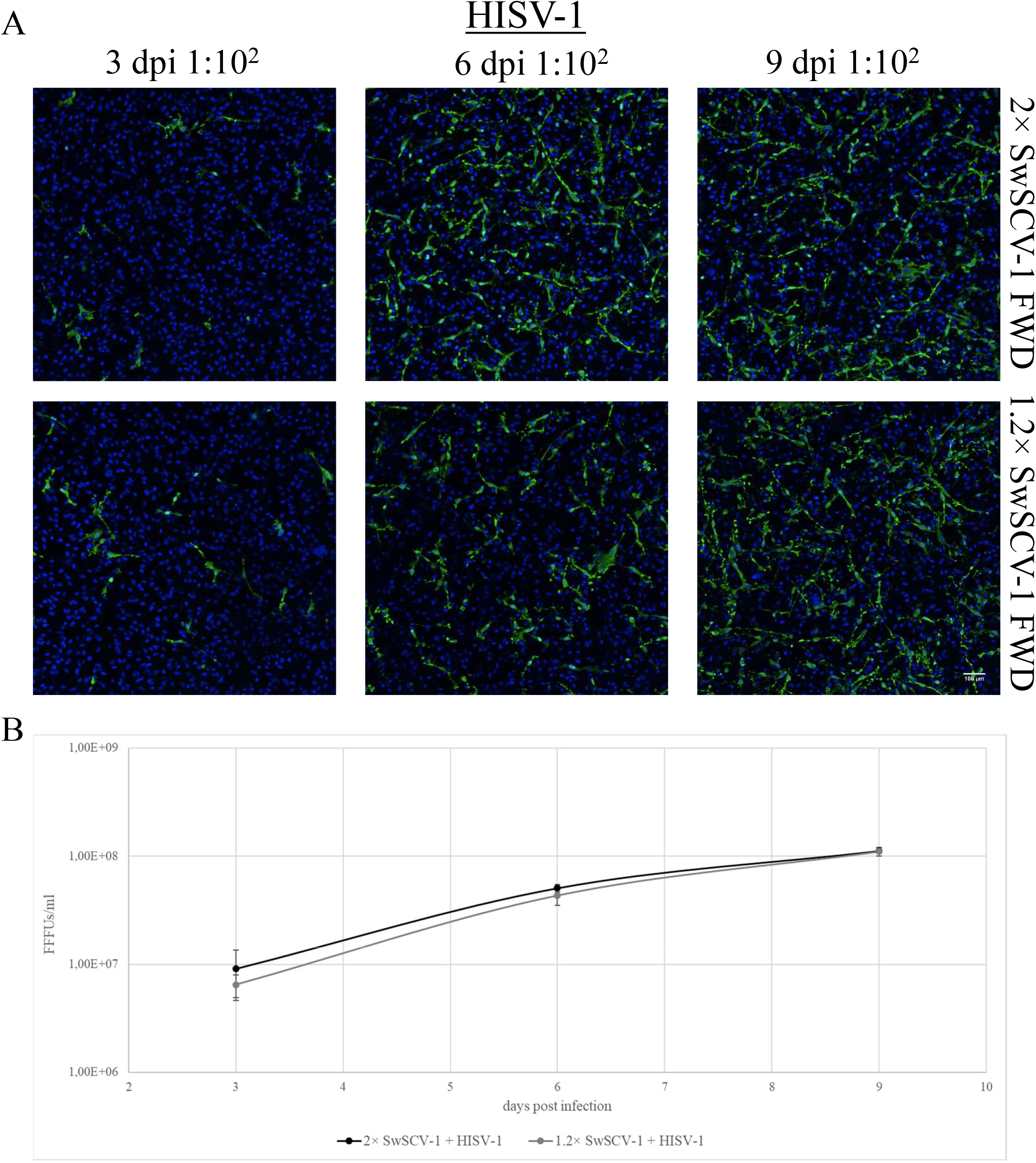
Superinfection of 2× and 1.2× SwSCV-1 FWD transfected I/1Ki cells leads to infectious particle production. **A)** Supernatants collected at 3, 6 and 9 days post HISV-1 superinfection from I/1Ki cells - transfected with 2× and 1.2× SwSCV-1 FWD constructs two weeks earlier - were used to inoculate clean I/1Ki cells. At four days post inoculation, the cells were fixed and stained using rabbit α-SwSCV-1 DAg antiserum and Alexa Fluor 488 labelled donkey anti-rabbit secondary antibody. Hoechst 33342 served for staining the nuclei. The top panels show clean I/1Ki cells infected with 100-fold diluted supernatant originating from HISV-1 superinfected 2× SwSCV-1 FWD transfected cells and the bottom panels with supernatant originating from HISV-1 superinfected 1.2× SwSCV-1 FWD transfected cells. The images were captured using Opera Phenix High Content Screening System (PerkinElmer) with 20× objective. **B)** Opera Phenix High Content Screening System (PerkinElmer) served to count the number of infected cells in **A**, which enabled the quantification of infectious particles per ml of growth medium in terms of fluorescent focus forming units (FFFUs – displayed on y-axis). The error bars represent standard deviation.

### 1.2× SwSCV-1 construct leads to persistent infection of the transfected cells

In our previous study, we showed that by maintaining I/1Ki cells after transfection with the 2× SwSCV-1 FWD construct, we could generate persistently SwSCV-1 infected cell lines (17). At the time of preparing this manuscript, we have maintained the I/1Ki-2×Δ cell line for 2.5 years, and IF staining for DAg shows the cell line to be persistently SwSCV-1 infected (Figure 6A). To compare the replication behavior of the shorter construct further, we transfected I/1Ki cells with 1.2× SwSCV-1 FWD and continued passaging the cells. Analysis of the cells by IF staining for DAg at 8 months post initial transfection indicates that also the 1.2× SwSCV-1 FWD construct can induce persistent infection in I/1Ki cells (Figure 6A). We compared the generated cell line, I/1Ki-1.2×Δ, to I/1Ki-2×Δ cells further by analyzing the amount of DAg expression using WB. The results show that DAg expression by I/1Ki-1.2×Δ cells is at least at the level observed in I/1Ki-2×Δ cells (Figure 6B), supporting the observation of a similar replication efficiency. To further compare the cell lines, we set up a near-infrared fluorescent northern blot assay for detection of the genomic RNA, antigenomic RNA, and SDAg mRNA. Northern blot of RNA isolated from I/1Ki-1.2×Δ and I/1Ki-2×Δ cells using a probe targeting the genomic RNA resulted in detection of a doublet band migrating at around 2.8 kilonucleotides (knt) from both cell lines (Figure 6C). To our surprise, the migration of the genomic RNA as compared to the RNA marker did not correspond to the expected 1.7-knt SwSCV-1 genomic RNA. Using higher RNA loading we detected a band migrating at around 5 knt, which likely represents the trimeric genome reported to be present in the infected cells by other researchers (43). We assume that the 2.8-knt band observed in SwSCV-1 infected cells represents the circular genome and speculate that the aberrant migration is due to the rod-like structure of the genomic RNA. We analyzed the persistently infected cells also for the presence of antigenomic RNA and succeeded in the detection when roughly 7 times more RNA was applied for the initial separation. As compared to the RNA marker, the bands detected using the antigenomic probe were approximately 3 knt by mobility (Figure 6C). In HDV the genomic RNA is approximately 5-10 times more abundant (44), and we think that, in addition to the rod-like structure, the high amount of genomic RNA further distorted the migration of the antigenomic RNA. We were unable to detect SDAg mRNA in the persistently infected cells, but detection of the antigenomic RNA by the probe (Figure 6C) suggests that the mRNA amount was below our detection limit.

**Figure 6.**
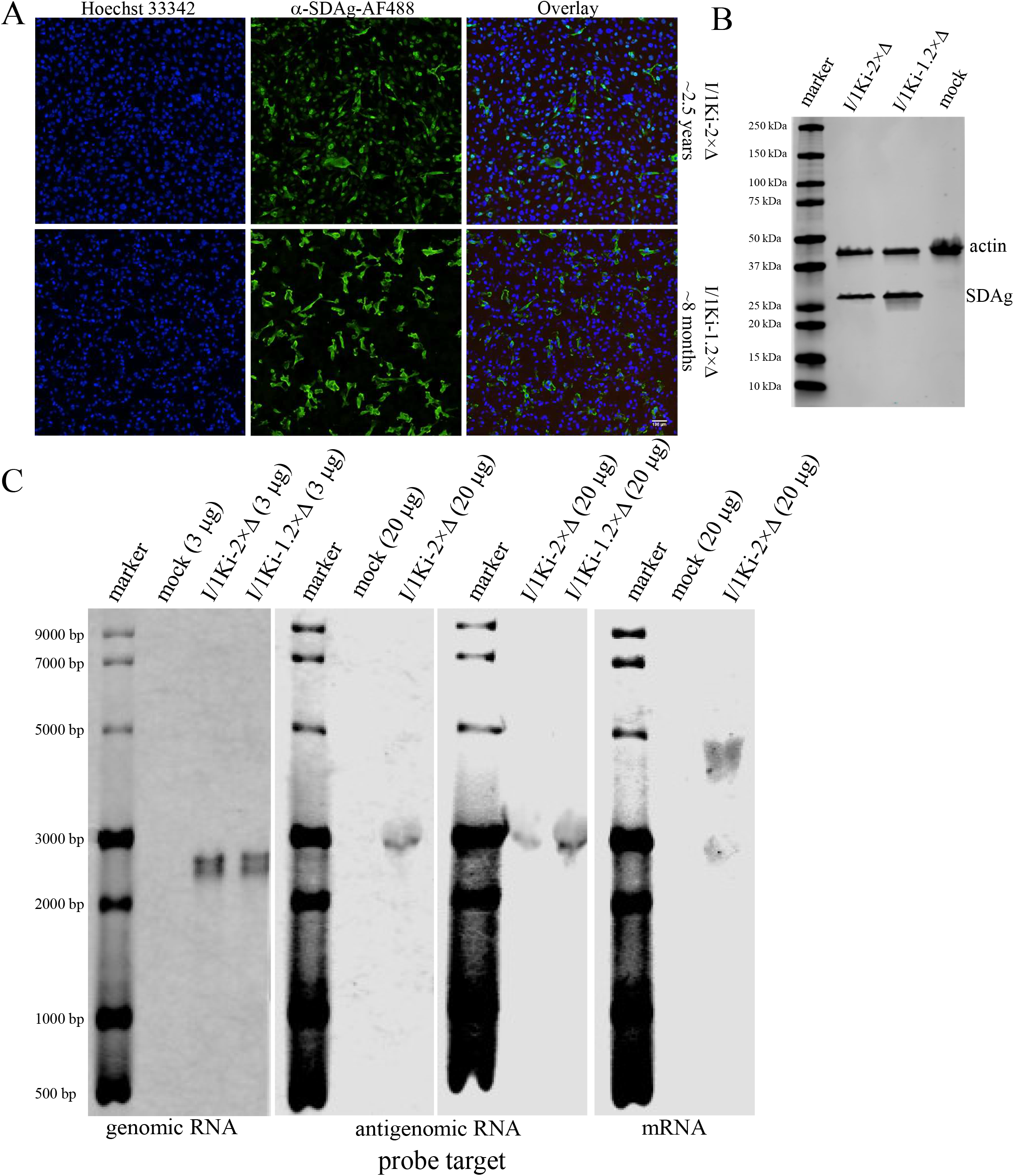
Comparison of persistently SwSCV-1 infected I/1Ki cells generated following transfection with 2× and 1.2× SwSCV-1 FWD constructs by immunofluorescence, western and northern blot. The 2×SwSCV-1 (I/1Ki-2×Δ) cell line was analyzed at approximately 2.5 years and 1.2×SwSCV-1 (I/1Ki-1.2×Δ) at approximately 8 months after initial transfection, during which the cell lines were passaged at 1-2 week interval. **A)** Rabbit α-SwSCV-1 DAg antiserum and Alexa Fluor 488 labelled donkey anti-rabbit secondary antibody served for IF staining of the fixed cells, and Hoechst 33342 for staining the nuclei. The top panels show staining of I/1Ki-2×Δ cells and the bottom panels the staining of I/1Ki-1.2×Δ cells. The left panels show staining of nuclei in blue, the middle panels show DAg staining in green, and the right panels show an overlay. The images were captured using Opera Phenix High Content Screening System (PerkinElmer) with 20× objective. **B)** Samples of I/1Ki-2×Δ cells and I/1Ki-1.2×Δ cells were separated on 4–20% Mini-PROTEAN TGX gels (Bio-Rad), transferred onto nitrocellulose, and the membranes probed with rabbit α-SwSCV-1 DAg antiserum and mouse monoclonal anti-pan actin antibody. The results were recorded using Odyssey Infrared Imaging System (LI-COR Biosciences). **C)** Indicated amounts of total RNA isolated from I/1Ki-2×Δ cells and I/1Ki-1.2×Δ cells, and clean I/1Ki were separated on agarose gel and transferred onto nylon membrane. Probes were targeting SwSCV-1 genomic (left panel), antigenomic RNA (middle panels), SDAg mRNA (right panel), and the bands of the marker served for visualizing the RNA targets. The results were recorded using Odyssey Infrared Imaging System (LI-COR Biosciences).

## DISCUSSION

The identification of novel deltaviruses significantly divergent from HDV (14, 15, 18-21), which until 2018 was the sole representative of the previously unassigned genus *Deltavirus*, has increased the interest in deltavirus research. The identification of novel deltaviruses in various host species without traces of hepadnaviruses by others and us led to questioning the strict association of HDV and HBV. Co-incidentally, Perez-Vargas and colleagues showed that HDV is able to use helper viruses other than HBV to form infectious particles (16). We demonstrated that SwSCV-1 efficiently utilized reptarena- and hartmaniviruses as its helpers, and that the co-expression of different arena- and orthohantavirus glycoproteins can drive infectious particle formation (17). Construction of infectious clones is the first step in demonstrating that the sequences recovered through metatranscriptomic analyses are indeed complete and capable of driving replication. We reported generation of such a clone by inserting two copies of the SwSCV-1 genome in head-to-tail fashion into a mammalian expression vector, pCAGGS (17). The same approach was hence proven functional for TSRV-1 (19) as well as *Taeniopygia guttata* deltavirus and *Marmota monax* deltavirus (21). The first HDV infectious clone contained a trimeric HDV genome, and the authors utilized a dimeric genome-containing plasmid with deletion in the DAg ORF to demonstrate the protein’s role in replication (4). Constructs containing multiples of the genome make synthetic inserts longer and complicate mutational studies because each modification needs to be inserted/generated multiple times. This motivated us to attempt generation of a 1.2× genome infectious clones for initiation of deltavirus replication. The availability of tools and reagents at hand forced us to focus on comparing the replication initiation between 2× and 1.2× SwSCV-1 genome clones in depth, but we were also able to demonstrate that a similar approach might work for the recently identified deltaviruses.

The replication of HDV occurs via rolling circle replication by cellular RNA polymerases (11) during which the genomic and antigenomic ribozymes cut the produced genome multimers into unit-length pieces (45). The recently identified deltaviruses presumably share the same replication strategy and possess the genomic and antigenomic ribozymes (14, 15, 19, 21, 46). Based on the HDV literature (47-49), we reasoned that duplicating the genomic and antigenomic ribozyme sequences would facilitate initiation of replication and/or production of unit-length genome (and antigenome). Indeed, RNA transfection studies with HDV have shown 1.2× genome copies to be most efficient in induction of virus replication (29), and a similar approach has been applied to generate HDV infectious clones (50). By applying the same principle, we constructed 1.2× genome infectious clones for HDV-1, SwSCV-1, TSRV-1, DabDV-1, and CITV-1 in both genomic and antigenomic sense, and tested the clones in I/1Ki cells, which efficiently support replication of SwSCV-1 following transfection with the 2× genome clone (17). In the REV constructs, the CAG promoter of the pCAGGS vector should mediate DAg translation, which provides a source of DAg for the first rounds of replication. For HDV, the reports suggest existence of an internal promoter that could drive the production of DAg (28, 51), such a promoter would presumably act in both our REV and FWD constructs and could also contribute to replication initiation. Indeed, the comparison of DAg production following transfection with 1.2× and 2× genome SwSCV-1 FWD and REV constructs demonstrated detectable DAg levels to appear earlier in cells transfected with REV constructs (Figure 4A). We thus used the IF staining of DAg from I/1Ki cells transfected with the 1.2× genome REV constructs (HDV-1, TSRV-1, DabDV-1, and CITV-1) to estimate the cross-reactivity of the anti-SwSCV-1 DAg antiserum (15) with DAg of the different viruses. The antiserum appeared to cross-react best with TSRV-1 DAg that is the closest relative of SwSCV-1 from the viruses included (19). The fact that HDV-1 DAg showed prominent nuclear staining, as would be expected based on the HDV literature, increased our confidence in the specificity of the IF staining. The antiserum appeared to cross-react moderately well with the DAg of DabDV-1, showing mainly cytoplasmic staining. While the CITV-1 DAg appeared to be barely detectable with the antiserum, the results did suggest production of CITV-1 DAg. Although the expression of DAg could come from internal promoters, we think that the DAg produced following transfection of FWD constructs is due to initiation of replication. While the inability of our SwSCV-1 DAg antiserum to cross-react with the DAgs of the other viruses tested likely explains the lower signal, it is also likely that the different deltaviruses are not replicating optimally in the *B. constrictor* cells. We hypothesize that each of the deltaviruses would be through promoter usage adapted to replicating in a specific host species.

The 1.2× genome construct design described herein could help to reduce the complexity of introducing mutations, and facilitate synthetic gene design for molecular biology studies of the recently identified deltaviruses and those that will be identified in the future. Our results with SwSCV-1 show that the 1.2× genome clone is at least as efficient as the 2× genome clone in initiation of replication. Furthermore, our results indicate that introduction of the insert in either genomic or antigenomic orientation functions equally well in the *B. constrictor* kidney cell model. Further studies with HDV-1, TSRV-1, DabDV-1, and CITV-1 in cell lines of various species could serve to demonstrate species specificity of the viruses, and to provide first evidence on the potential role of deltavirus promoters in mediating species-specific replication.

## ACKNOWLEDGEMENTS

The authors wish to acknowledge Dr. Antti Hassinen of the FIMM (Institute for Molecular Medicine Finland) High Content Imaging and Analysis (FIMM-HCA) for expert help in imaging and quantification of the IF staining for titration. The study was funded by the Academy of Finland (to JH, grant numbers: 308613, 314119, and 335762), the funding body had no role in the study design, interpretation of the results or preparation of the manuscript.

